# Loss of CIB2 causes non-canonical autophagy deficits and visual impairment

**DOI:** 10.1101/2020.09.18.302174

**Authors:** Saumil Sethna, Patrick A. Scott, Arnaud P.J. Giese, Todd Duncan, T. Michael Redmond, Saima Riazuddin, Zubair M. Ahmed

## Abstract

Non-canonical autophagy or LC3-associated phagocytosis (LAP) is essential for the maintenance and functioning of the retinal pigment epithelium (RPE) and photoreceptors. Although molecular mechanisms still remain elusive, deficits in LAP have been found to be associated with age-related retinal pathology in both mice and humans. In this study, we found that calcium and integrin-binding protein 2 (CIB2) regulates LAP in the RPE. Mice lacking CIB2, both globally and specifically within RPE, have an impaired ability to process the engulfed photoreceptor outer segments due to reduced lysosomal capacity, which leads to marked accumulation of improperly digested remnants, lipid droplets, fused phago-melanosomes in RPE, and impaired visual function. In aged mice, we also found marked accumulation of drusen markers APOE, C3, and Aβ, along with esterified cholesterol. Intriguingly, we were able to transiently rescue the photoreceptor function in *Cib2* mutant mice by exogenous retinoid delivery. Our study links LAP and phagocytic clearance with CIB2, and their relevance to the sense of sight.

## INTRODUCTION

Highly specialized polarized neurons, known as rod and cone photoreceptors (PR), serve as the site of transduction where irradiant light is converted into neuronal signals in the eye. PR outer segments (OS) are dedicated to phototransduction, however they depend heavily on support from the retinal pigment epithelium (RPE), a single-layer of polarized cobblestone-shaped post-mitotic cells. RPE has multiple functions essential to normal vision, such as absorbing excess light, regenerating vitamin A-derived chromophores, and shuttling nutrients from the sub-retinal space to the PR and metabolites^1^. RPE also plays a unique but integral role in PR survival and function. It engulfs the shed OS discs, and subsequently, obtain clearance via LC3-associated phagocytosis (LAP) ^2^. The OS only grow from one end, but during the process of OS renewal, the distal-most photo-damaged disks are pruned by the RPE in a diurnal light-entrained circadian manner. An initial burst of OS binding and ingestion occurs immediately after light onset, followed by clearance through RPE lysosomal machinery ^2-12^. Prompt and efficient OS engulfment and the subsequent phagolysosomal clearance are both essential to help prevent the age-related buildup of undigested debris within and below the RPE. Furthermore, for all practical purposes, the PR and RPE act as a single functional unit, with dysfunction of one leading to dysfunction of the other.

Growing evidence suggests that RPE dysfunction, specifically in OS digestion and clearance, leads to age-related photoreceptor dysfunction without gross retinal degeneration ^2,9,13^. For instance, RPE-specific deletion of ATG5, caveolin-1, or the β-crystallin protein leads to the impaired digestion of OS but does not cause gross retinal degeneration. However, the PR function (measured by electroretinography (ERG)) is substantially impaired ^2,9,13^. Despite the significance of RPE-mediated OS processing, the precise molecular parameters of phagolysosomal digestion remain poorly understood. Further, phagocytosis/ autophagy defects have been implicated in the dry form of age-related macular degeneration (AMD) ^14^. Therefore, deciphering the molecular regulators has significant ramifications for understanding the fundamental retinal aging process as well as the etiology of certain disorders.

We previously identified that CIB2 impairment was associated with deafness and/or vision deficits in humans, zebrafish, and drosophila ^15^. Down-regulation of *cib2* in drosophila significantly reduces photoresponse amplitudes and causes light-dependent retinal degeneration ^15^. However, the molecular mechanism of such CIB2-associated vision loss remains unknown. Here, we show that the ablation of CIB2, specifically in the RPE, causes an age-related phenotype in mice and dysregulation of phagolysosomal processing of OS. We reason that abundant accumulation of large lipid-filled vacuoles and sub-retinal deposits might impede transport of retinoids back to the PRs. This in turn leads to reduced ERG amplitudes, even without gross PR damage. Supporting this idea, we found that providing exogenous retinoids to two different *Cib2* mouse models improves their ERGs in two different treatment paradigms. Finally, our study establishes a proof-of-concept for the use of retinoids to treat human ocular diseases, especially age-related RPE pathologies.

## RESULTS

### Loss of CIB2 leads to age-related PR dysfunction and RPE pathology

Within the retina, CIB2 is expressed in the RPE, PRs, and certain ganglion cells (**Supplementary Fig. 1a-e**). To determine the exact function of CIB2 in the mammalian retina, we used *Cib2*^*tm1a*^ mutant mice (**Fig. 1a**, *Cib2*^*KO*^ henceforth) ^16^ that lack CIB2 in the RPE (**Supplementary Fig. 1f**) as well as other retinal layers. Non-invasive *in vivo* scotopic full-field electroretinograms (ERGs, illustrated in **Fig. 1b**), which preferentially analyze rod PR function, revealed no difference in a-wave (derived primarily from the photoreceptor layer) or b-wave (derived from the inner retina, predominantly Müller and outer nuclear bipolar cells) amplitudes in one-month-old *Cib2* deficient mice. However, at three, six and nine months, both *Cib2*^*KO/+*^ and *Cib2*^*KO/KO*^ mice exhibited similar age-related declines (∼20-30%) in both a- and b-wave amplitudes, as compared with those of wild type (WT) mice (**Fig. 1c**). However, the declines were not demonstrated in latency or oscillatory potential, suggesting the inner retinal function was not impacted (**Supplementary Fig. 2a, c**. In contrast, the b-wave amplitude of the photopic ERG, which evaluates cone PRs, was similar across all three genotypes (**Supplementary Fig. 2b**). These results suggest that both haploinsufficiency and complete loss of CIB2 leads to rod PR dysfunction in mice.

**Figure 1:**
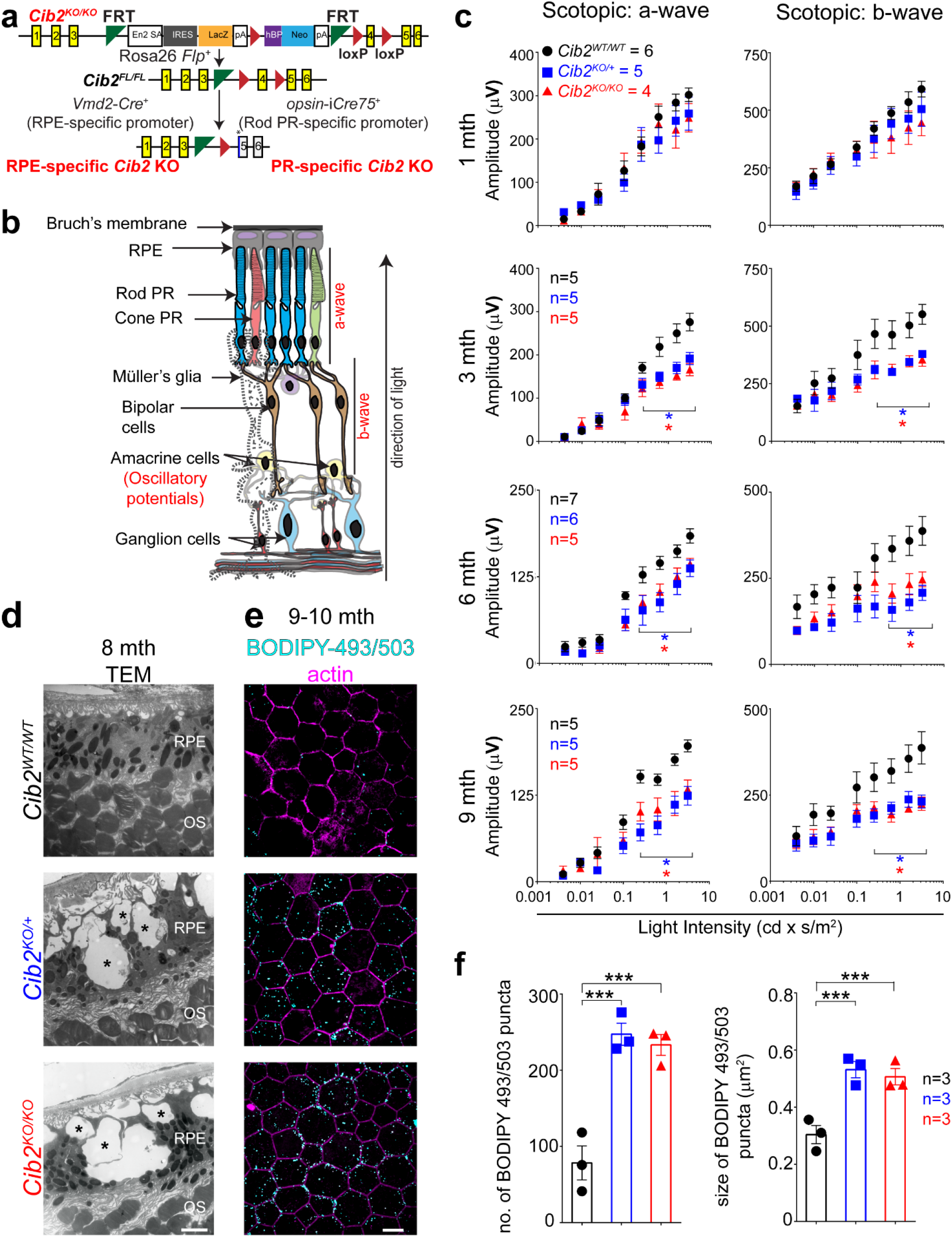
Loss of CIB2 leads to age-related photoreceptor (PR) dysfunction and RPE pathophysiology. **a** Schematic of specific constructs used to generate mouse models. **b** Schematic representation of the multiple retinal layers. ERG a-wave originates from photoreceptors (PR), while b-wave amplitude originates from Müller glia and outer nuclear bipolar cells, and oscillatory potential comes from the amacrine cells. RPE – retinal pigment epithelium. **c** Quantification of scotopic responses from WT (*Cib2*^*WT/WT*^), *Cib2*^*KO/+*^, and *Cib2*^*KO/KO*^ mice at indicated ages reveals progressive loss of both a- (*left* panels) and b-wave (*right* panels) amplitudes in *Cib2* deficient mice. **d** TEM micrographs of RPE/ outer segment (OS) interface from eight-month-old mice revealed vacuoles (*) and basal infoldings loss specifically in *Cib2*^*KO/+*^ and *Cib2*^*KO/KO*^ mutants (n = 3 per genotype). *Scale bar*, 2 μm. **e** Representative photomicrographs of RPE whole mounts from nine-to-ten month old mice for denoted genotype showing accumulation of neutral lipid marker BODIPY-493/503 (*cyan*) in *Cib2* deficient mice. Phalloidin (*magenta*) was used to decorate actin cytoskeleton. *Scale bar*, 10 µm. **f** Quantification of average number (*left*) or average size (*right*) of BODIPY-493/503-stained puncta per mouse shown in **(e)**. At least three images per mouse (∼25 cells/image) were quantified and averages per mouse are shown for WT, *Cib2*^*KO/+*^, and *Cib2*^*KO/KO*^ mice (*n* = 3 each). Data presented as mean±SEM; each *point* represents an individual animal. One-way ANOVA and Bonferroni *post hoc* test, *p* value of <0.05 (*) or <0.001 (***).

To determine if corresponding anatomical changes occurred with progressive loss of PR function, we evaluated the morphology of the retina in two and eight-to nine-month-old *Cib2*-deficient mice. Light microscopy-based morphometric analysis of the specific retinal strata revealed no significant differences in *Cib2*^*KO/+*^ and *Cib2*^*KO/KO*^ compared to those of WT mice at either age (**Supplementary Fig. 2d, e**). However, at the transmission electron microscopic (TEM) level, we observed vacuoles with undigested membranous material and loss of basal infoldings in the RPE of *Cib2*^*KO/+*^ and *Cib2*^*KO/KO*^ mice, but not in age-matched WT mice (**Fig. 1d**). Staining with the neutral lipid tracer BODIPY™ 493/503 revealed two-fold more lipid droplets in the RPE of aged *Cib2*^*KO/+*^ and *Cib2*^*KO/KO*^ mice (**Fig. 1e, f**). Together, these results suggest that loss of CIB2 causes excessive lipid accumulation in RPE and is associated with progressive rod PR dysfunction.

### Selective ablation of CIB2 in RPE but not in PRs recapitulates retinal pathophysiology

In the mammalian retina, PRs and RPE are codependent, therefore, the functional impairment of one layer negatively impacts the other ^2,9,17,18^. To pinpoint the anatomical site of age-related rod PR dysfunction seen in global *Cib2*-deficient mice, we generated two cell type-specific mutant strains, in which *Cib2* expression was ablated either in rod PR (designated PR-*Cre+*) using the rod PR-specific *rhodopsin*–*iCre75* ^19^, or in the RPE (designated RPE-*Cre+*) using the RPE-specific *Vmd2* promoter ^20^ (**Fig. 1a** and **Methods**). ERG analyses revealed that the RPE-specific *Cib2*^*KO*^ (*Cib2*^*flox/flox*^; RPE-*Cre+* and *Cib2*^*flox/+*^; RPE-*Cre+*) mice had reduced ERG amplitudes compared to the control mice (*Cib2*^*+/+*^; RPE-*Cre+*) as early as 3 months of age. Similar to *Cib2* global knockout mice, this declined even further with age (**Fig. 2a**). In contrast, we found no differences in either a- or b-wave amplitudes with PR-specific *Cib2*^*KO*^ (*Cib2*^*flox/flox*^; PR-*Cre+* and *Cib2*^*flox/+*^; PR-*Cre+*) mice compared to control mice at any tested age (**Fig. 2b; Supplementary Fig. 3a, b**). Holistically analyzed, the retinal functional and morphological analyses in three mutant strains suggests that rod PR dysfunction occurs secondary to the loss of CIB2 function in RPE but not in rod PRs.

**Figure 2:**
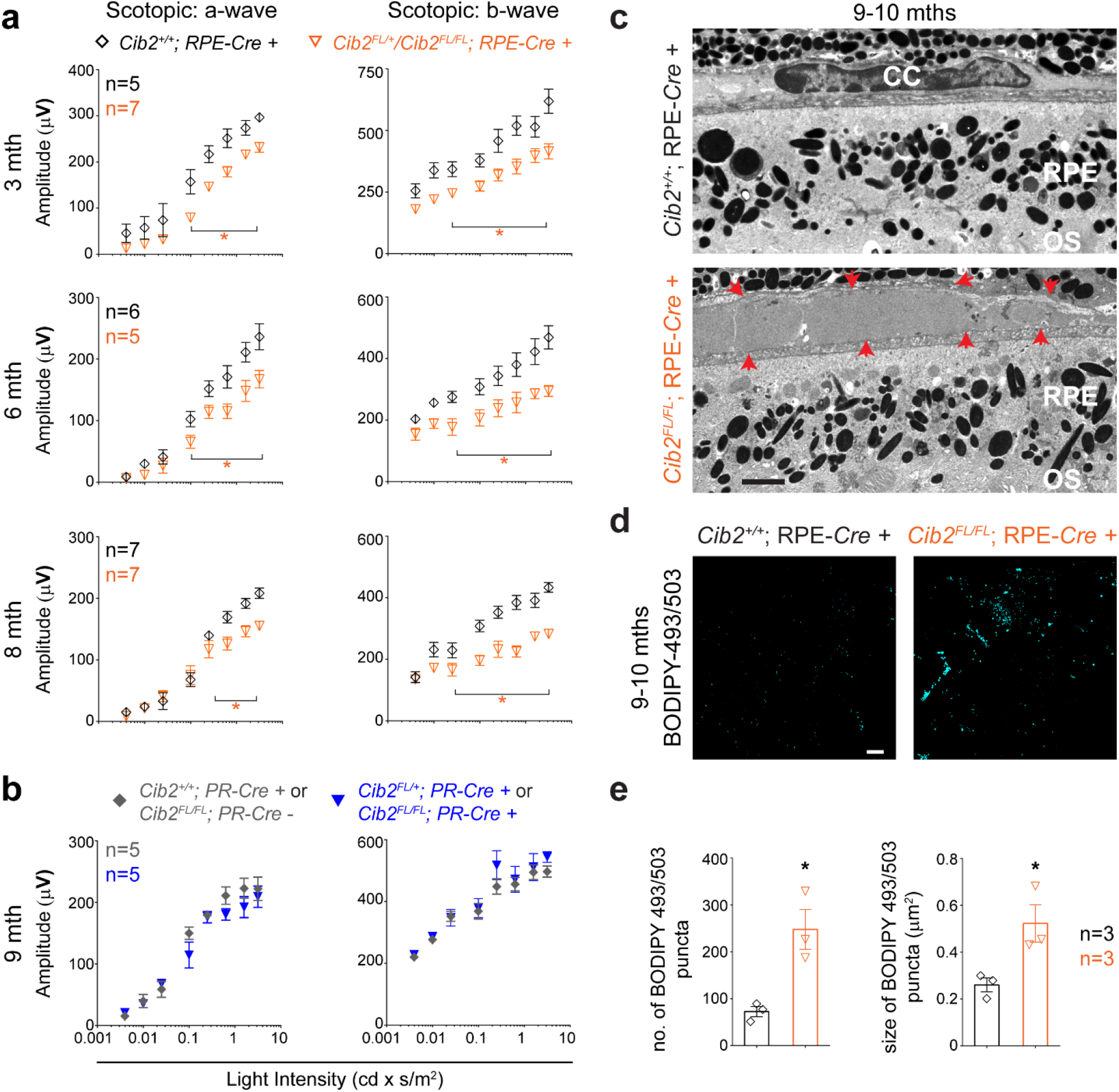
Loss of CIB2 specifically from RPE, but not rod photoreceptors recapitulates age-related phenotype. **a, b** Quantification of scotopic a- (*left* panels) and b-wave (*right* panels) amplitudes for RPE-*Cre+* (**a)** and PR-*Cre* mice (**b**) at indicated ages. Loss of CIB2 specifically from RPE but not rod photoreceptors (PR) resulted in ERG deficits. **c** TEM micrographs of nine-to-ten month old RPE-*Cre+* control *Cib2*^*+/+*^ (*top* panel) and *Cib2*^*FL/FL*^ (*bottom* panel) mice (n=3 per genotype). Red arrows (*bottom* panel) indicate boundaries of deposits present between RPE and choroid tissue in RPE-specific mutant mice. *Scale bar*, 2 μm. RPE – retinal pigment epithelium; OS – outer segment; CC – choriocapillaris. **d** Representative photomicrographs of RPE whole mounts from nine-to-ten month old mice for denoted genotype demonstrate accumulation of neutral lipids in RPE-specific *Cib2* mutant mice. *Scale bar*, 10 µm. **e** Quantification of average number (*left*) or size (*right*) of BODIPY-493/503 puncta per mouse shown in **(d)**. At least three images per mouse (∼25 cells/image) were quantified and averages per mouse are shown for denoted genotypes (n = 3 per genotype). Data presented as mean±SEM. Each data point represents an individual mouse. Student unpaired t-test, *p* value <0.05 (*).

With respect to the thickness of RPE- and PR-specific *Cib2*^*KO*^ mice retinal strata, we found no morphometric differences (**Supplementary Fig. 3c, d**, data not shown), however, TEM revealed sub-RPE deposits in RPE-specific *Cib2*^*KO*^ mice (**Fig. 2c**) that were absent in PR-specific mutants (**Supplementary Fig. 3e**). As seen in *Cib2*^*KO*^ global mice, we also found the presence of an increased amount of lipid droplets in the RPE of RPE-specific *Cib2*^*KO*^ mice (**Fig. 2d, e**). We further confirmed the presence of these lipid droplets in nine-to ten-month old RPE-specific mutant mice by staining frozen transverse sections with Oil Red O (**Fig 3a**). Lastly, we also found increased filipin staining specifically under RPE, which revealed the accumulation of both unesterified and esterified cholesterol (**Fig 3b**). Esterified cholesterol, in particular, has been shown to accumulate in drusen and deposits in the Bruch’s membrane in humans suffering with dry AMD ^21^.

**Figure 3:**
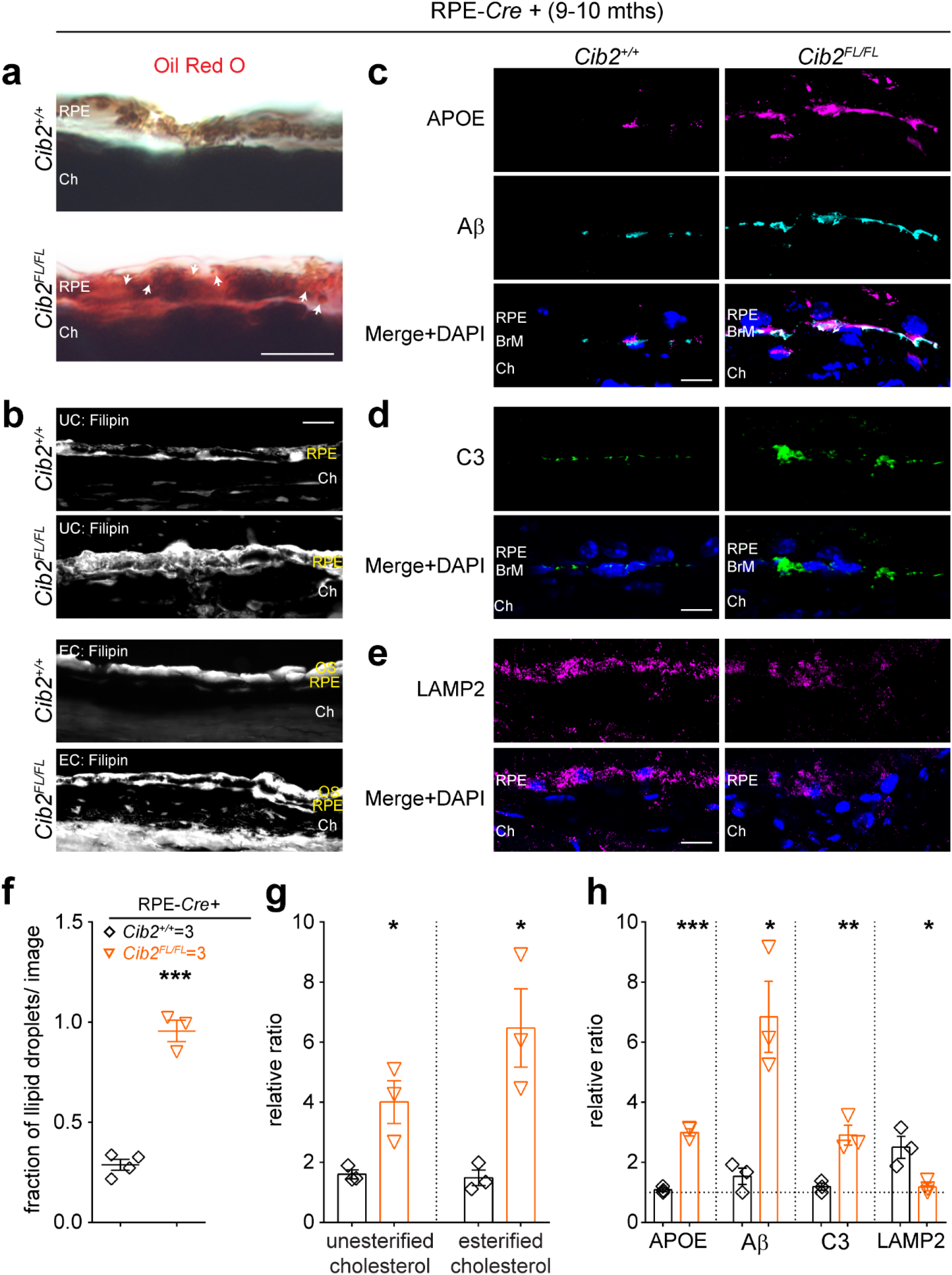
*Cib2* mutant mice mimic the drusen pathologies reported in human RPE suffering with dry AMD. **a** Oil red O staining shows a marked accumulation of lipids within and below the RPE in mutant mice. Arrows indicate lipid droplets. **b** Filipin staining for unesterified cholesterol (UC, *top* two panels) and esterified cholesterol (EC, *bottom* two panels) shows marked accumulation of both UC and EC, particularly in mutant mice, below the RPE. **c, d** Confocal micrographs of cryosections of nine-to-ten month old mice of denoted genotype shows the accumulation of specific proteins APOE, Aβ **(a)**, and C3 **(b)** below the RPE in mutant mice. **e** Confocal micrographs of cryosections of nine-to-ten month old mice of denoted genotype show that the levels of lysosomal protein LAMP2 are lower within the RPE of mutant mice. *Scale bar*, 10 μm (**a, c**), 20 μm (**d, e**). RPE – retinal pigment epithelium; BrM – Bruch’s membrane; Ch – choroid; OS – outer segments. **f** Quantification for fraction of lipid droplets for images presented in **a. g** quantification of relative ratio of denoted lipid species for images shown in **b. h** quantification of relative ratio of denoted proteins levels for images shown in **c-e**. At least three images per mouse were quantified and averages per mouse are shown for denoted genotypes (n = 3 per genotype). Data presented as mean±SEM. Each data point represents an individual mouse. Student unpaired t-test, *p* value of <0.05 (*), <0.01 (**), or <0.001 (***).

### Molecular characterization of sub-RPE deposits in *Cib2*-deficient mice

As presented in humans, AMD is a progressive, age-related, and degenerative disease of the retina. AMD is categorized as either wet or dry, due to either choroidal neovascularization or progressive thinning of macula layers, respectively ^22^. Sub-RPE drusen are a prominent feature observed in humans suffering with dry AMD ^23^. Protein-level analysis of drusen found in dry AMD patients, revealed accumulation of proteins such as apolipoprotein E (APOE) ^24^, β-Amyloid (aβ), and complement factor 3 (C3), a genetically validated AMD risk factor ^25-27^. Meanwhile, a recent study showcased that the levels of LAMP2/CD107b (a lysosomal membrane protein) are reduced in AMD RPE/choroid tissues ^28^. Observation of similar sub-RPE deposits in TEM micrographs in *Cib2* mutant mice prompted us to investigate the molecular composition of the deposits. We evaluated and found that in nine-to-ten month old RPE-specific *Cib2*^*KO*^ mice there was marked accumulation of APOE, β-Amyloid (aβ), and C3 (**Fig 3c, d**) below the RPE and reduced expression of LAMP2/CD107b within the RPE (**Fig 3e**). Ultimately, our data shows that *bona fide* AMD drusen markers and lipid species are enriched in the sub-RPE deposits in CIB2 mutant mice, which suggested a role of CIB2 in human AMD ^29^.

### Loss of CIB2 leads to dysregulated clearance of photoreceptor outer segments

Defects in the OS renewal process in mice can lead to the formation of vacuoles and an accumulation of lipids ^9^, a phenotype similar to that presented in *Cib2*^*KO*^ mice. We reasoned that a lack of CIB2 may lead to debris and vacuole formation in older mice (8-10 months) due to faulty phagolysosomal processing of the OS by the RPE early in life (2-4 months). We took advantage of the characteristic diurnal circadian nature of RPE-mediated OS phagocytosis to quantify engulfment (early after light onset) and digestion (rest of the day) by *in situ* quantification of phagosomes within the RPE, at specific times after light onset. The OS are composed of 50% lipids and 50% proteins, and rhodopsin constitutes about 80% of rod OS protein content. Thus, any rhodopsin found in the RPE is from phagocytosed OS and can be used as a marker to track the OS renewal process ^8,18^. *In situ*, we found that counts of opsin-phagosomes in whole mounts of RPE/choroid one hr after light onset, corresponding to peak of OS ingestion, were similar in *Cib2*^*KO*^ and WT mice, indicating that binding and ingestion of OS were unaffected by *Cib2* deficiency (**Fig. 4a *top panel*, 4b**). However, *Cib2*^*KO*^ mice exhibited significantly higher numbers of opsin-phagosomes eight and twelve hours following the onset of light, representing slower phagolysosomal clearance (**Fig. 4a *bottom panel*, 4b; Supplementary Fig. 4a, b**). The average diameters of opsin-phagosomes decreased in tandem with time from onset of light-on in WT animals. On the other hand, the decreases in *Cib2*^*KO*^ mice were delayed, further confirming that *Cib2* deficiency leads to phagolysosomal dysfunction (**Supplementary Fig. 4c**). TEM imaging of two-month-old mice retinae that were dissected eight hours after light onset revealed that *Cib2*^*KO/+*^ and *Cib2*^*KO/KO*^ mice exhibited marked accumulation of improperly-digested remnants, lipid droplets fused with undigested material, and/or fused phago-melanosomes (**Fig. 4c, d**). Similar deficits were observed in RPE-specific *Cib2*^*KO*^ mice eight hours after light onset (**Fig. 4f, g**). Lastly, we also found sub-RPE deposits and debris in mutant mice but not in the retinae of WT mice (**Fig. e**).

**Figure 4:**
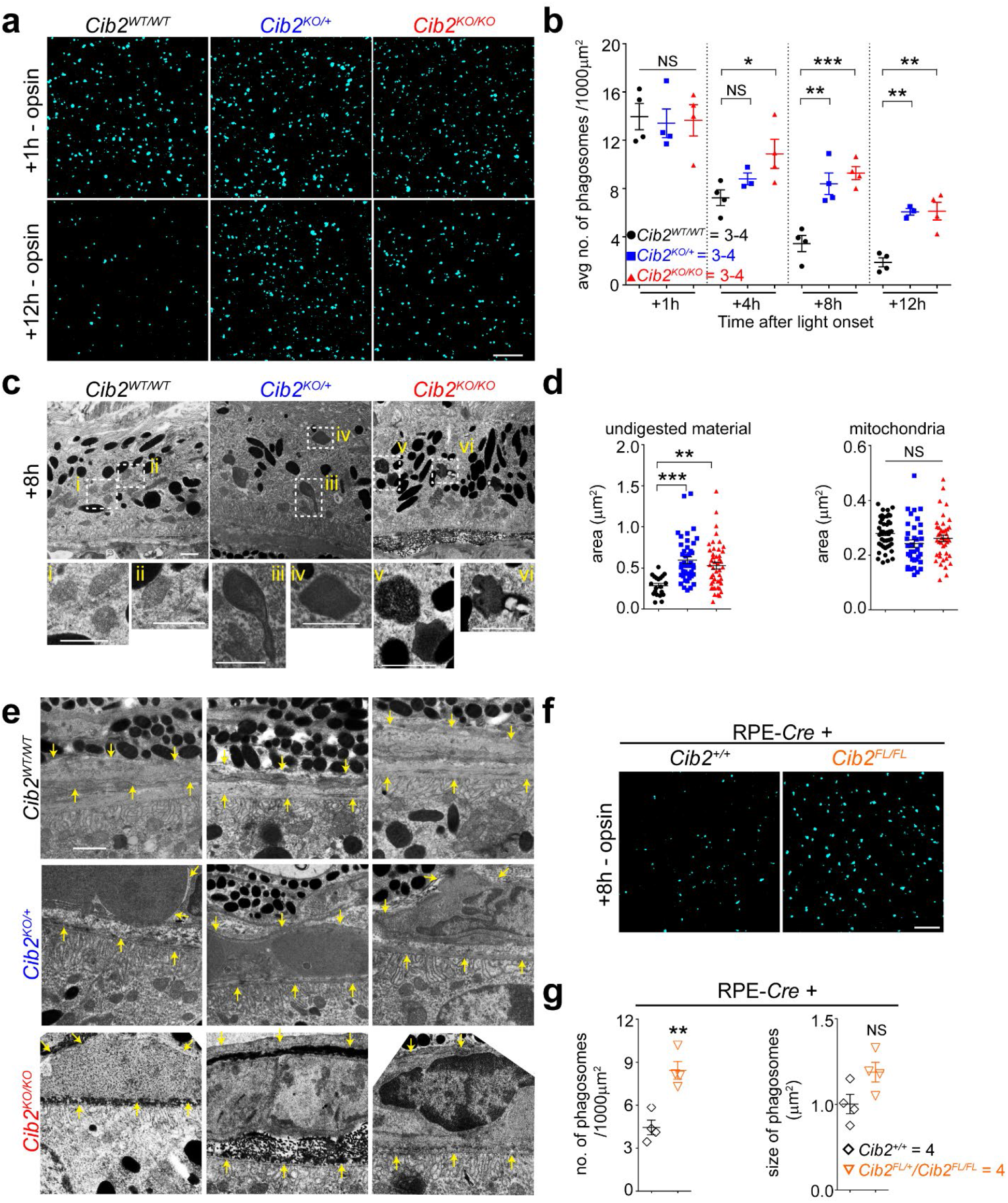
Deficiency of CIB2 resulted in impaired phagolysosomal processing of OS. **a** RPE whole mounts from three to four month old mice immunostained with opsin Ret-P1 antibody (*cyan*) shows initial binding and ingestion of opsin-phagosomes is similar across genotypes (*top* panel), however, the phagolysosomal digestion is slower in mutant mice (*bottom* panel) as compared to the WT control. *Scale bar*, 10 µm. **b** Quantification of opsin-phagosomes shown in panel **a and supplementary fig. 4a. c** TEM micrographs of two-month-old mice euthanized eight hours after light onset show accumulation of undigested material, and fused remnants of phago-melanosomes in mutant mice. **(i – vi)** *Inset*s show magnified images of normal phagosomes in WT mice and improperly processed phagosomes in only in *Cib2*^*KO/+*^ and *Cib2*^*KO/KO*^ mice. *Scale bar*, 1 μm. **d** Quantification of individual phagosomal (*left*) and mitochondrial (*right*) areas (as control) for images shown in panel **d** further confirm marked accumulation of phagosomal area. All phagosomes and 15 mitochondria /image in three to five images per mouse were counted (n=3 per genotype). **e** TEM micrographs from two month old mice show marked thickening and accumulation of debris under the RPE in mutant mice. Yellow arrows indicate boundary between basal surface of RPE and choroid. *Scale bar*, 1 μm. **f** Representative RPE whole mounts image from three-to-four month old mice immunostained with opsin Ret-P1 antibody (*cyan*) show slower phagolysosomal clearance, quantified in panel **g**, in RPE-specific *Cib2* mutant mice. *Scale bar*, 10 µm. At least three images per mouse (∼30 cells/ image) were quantified and average per animal are shown for denoted genotypes. Data presented as mean±SEM. Each data point represents an average per individual mouse or experiment unless specified. One-way ANOVA and Bonferroni *post hoc* test (**b, d**) or unpaired two-tailed *t* test (**g**), *p* value of <0.05 (*), <0.01 (**), or <0.001 (***), NS – not significant.

Proper clearance of ingested OS requires optimally functioning lysosomal machinery. In the RPE, the aspartyl protease cathepsin D is essential for phagosome digestion ^4,18,30,31^. Cathepsin D is translated as ∼50 kD pro-cathepsin D and matures to its active form (∼30 kDa) in the low-pH lysosomal milieu. RPE flat mounts *ex vivo* probed with BODIPY-pepstatin A, which binds specifically to mature cathepsin D, showed a marked fold reduction of about 2.5 in indirectly-stained lysosomes, and hence available mature cathepsin D in *Cib2*^*KO/+*^, *Cib2*^*KO/KO*^, and RPE-specific *Cib2*^*KO*^ mice (**Fig. 5a-d**, and **Supplementary Fig. 5a, 5b –** *left panel*), but not in PR-specific *Cib2*^*KO*^ mice (**Fig. 5e** and **Supplementary Fig. 5b –** *right panel*). Further, immunoblotting revealed markedly reduced levels of mature cathepsin D, consistent with the BODIPY-pepstatin A labeling, LAMP-1, and ATG5-ATG12 complex in *Cib2*^*KO/KO*^ mice, both one and twelve hours after light onset (**Fig. 5f, g**).

**Figure 5:**
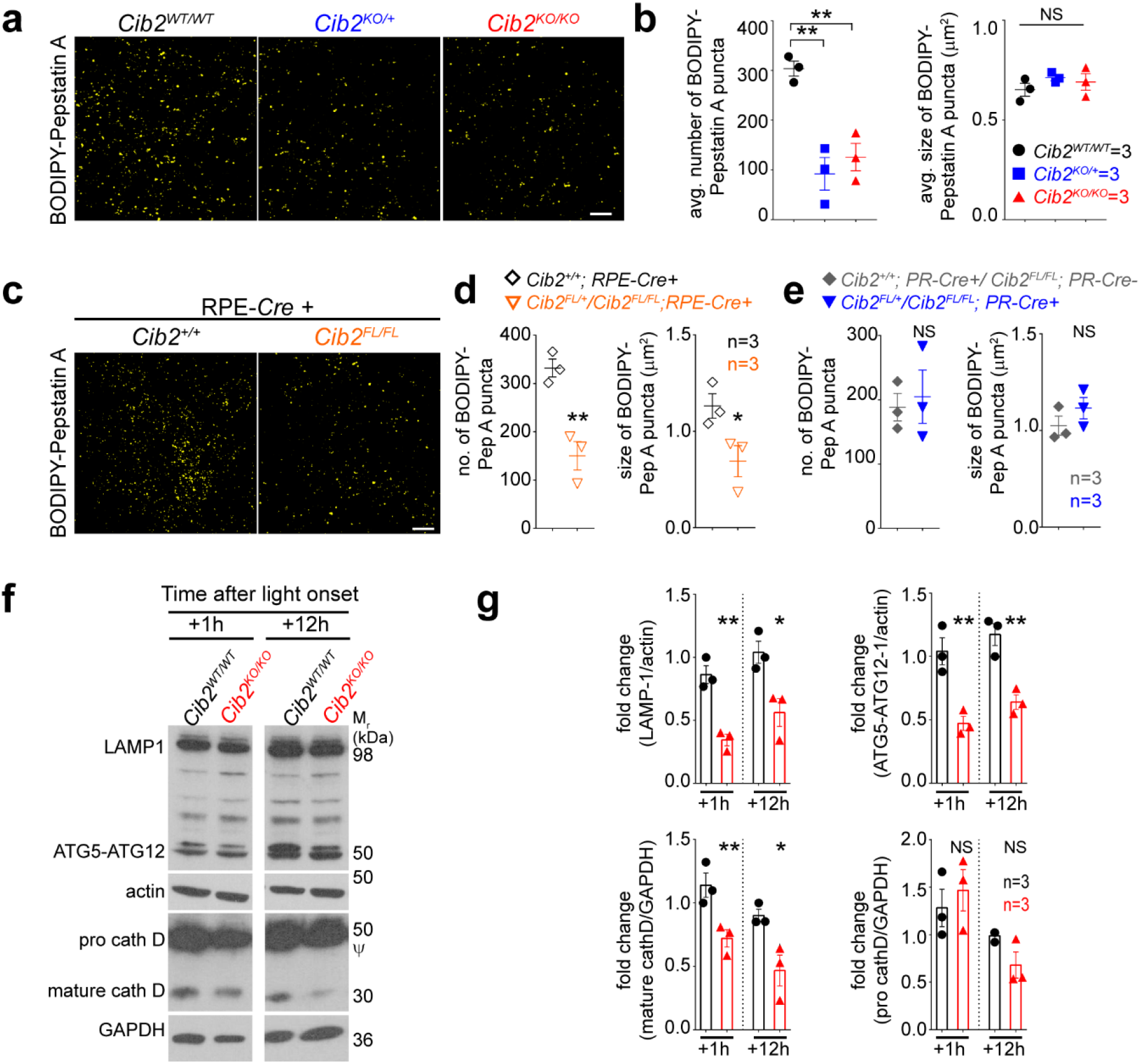
Deficiency of CIB2 resulted in impaired lysosomal machinery. **a** RPE whole mounts from three-to-four month old mice stained with BODIPY-pepstatin A, a dye which binds to active cathepsin D only, demonstrate fewer indirectly stained lysosomes, quantified in panel **b**, in *Cib2* deficient mice. RPE distinguished by phalloidin staining are shown in **supplementary figure 5a**. *Scale bar*, 10 µm. **c** Similarly, RPE whole mounts from three-to-four month old mice show that RPE-specific *Cib2* mutants have less BODIPY-pepstatin A stained lysosomes. RPE distinguished by phalloidin staining are shown in **supplementary figure 5b**. *Scale bar*, 10 µm. **d, e** Quantification of average numbers (*left*) or sizes (*right*) of BODIPY-pepstatin A puncta per mouse in RPE-*Cre+* (**d**) and PR-*Cre* (**e**) mice shown in panel **c** and **supplementary figure 5b. f, g** Representative immunoblots (**f**) for denoted proteins from RPE/choroid lysates obtained from mice euthanized either one or twelve hours after light onset for denoted genotypes indicates that lysosomal/autophagy proteins levels are decreased in mutant mice at both time points assessed. *ψ*, pro-cathepsin D immunoblot is overexposed for mature-cathepsin D to be observable, quantified in **g**. N = 3/genotype/time point. Pro-cathepsin D levels were quantified from non-saturated exposures. At least three images per mouse (∼30 cells/ image) were quantified and average per animal are shown for denoted genotypes. Data presented as mean±SEM. Each data point represents an average per individual mouse or experiment unless specified. One-way ANOVA and Bonferroni *post hoc* test (**b**) or unpaired two-tailed *t* test (**d, e, g**), *p* value of <0.05 (*) or <0.01 (**), NS – not significant.

Complementary to the findings of *Cib2* mutant mice, overexpressing CIB2 in RPE-J cells increased LAMP-1 and ATG5-ATG12 complex proteins levels (**Supplementary Fig. 5c, d**). To assess whether CIB2 overexpression has functional impact, we performed *in vitro* pulse-chase phagocytosis assays ^18^ followed by immunoblotting for opsin in RPE-J cells (schematic – **Supplementary Fig. 5e**, *left* panel). Again, in agreement with our findings in mice, we detected similar opsin levels bound initially (pulse-0 h) with control or CIB2 overexpression. However, 6 hours later, corresponding to the digestion phase, we found less opsin in cells with excessive CIB2, suggesting that CIB2 overexpression alone suffices to boost phagolysosomal digestion (**Supplementary Fig. 5e, f**). Collectively, our results suggest that lack of CIB2, specifically in the RPE, causes abnormal OS phagolysosomal processing due to reduced numbers of lysosomes and lysosomal protein levels, whereas transient CIB2 overexpression augments OS clearance.

### Rescue of PR function by evading the RPE-retinoid cycle

RPE is essential for the visual cycle regeneration of 11-*cis* retinal, the chromophore of photoreceptor opsins. Upon photon absorption, 11-*cis* retinal is photo-isomerized to all-*trans* retinal, thereby activating opsin and initiating the phototransduction cascade ^32,33^. All-*trans* retinal is regenerated to 11-*cis* retinal, which is then recycled back to the photoreceptors via a series of transport and enzymatic steps, occurring primarily in the RPE (illustrated in **Fig. 6a**). The ERG deficits without significant photoreceptor damage led us to investigate this key visual pathway. First, we quantified the absolute retinoid levels (11-*cis* oxime, all-*trans* oxime, retinyl esters, and A2E) within the retina or RPE/choroid in two-to-three month and eight-to-nine month old *Cib2*^*KO/+*^, *Cib2*^*KO/KO*^. However, no significant differences were observed when compared to control age-matched WT mice (**Supplementary Fig. 6a, b**). We reasoned that the vacuoles observed in the RPE may partially sequester retinoids, rendering them unavailable to re-enter the PR. Therefore, we tested whether we could rescue the ERG defects via exogenous chromophore, thus bypassing the RPE visual cycle altogether. For these studies, we used two different treatment paradigms. First, we injected aged global *Cib2*^*KO*^ (either heterozygous or homozygous) mice once with either 9-*cis* retinal (an analog of 11-*cis* retinal) or vehicle. Then, we assessed their visual function by ERG the following day, after dark adaptation. Treatment of *Cib2*^*KO*^ mice with 9-*cis* retinal restored their ERG amplitudes to near WT levels with significant differences in scotopic a- and b-wave amplitudes as compared to vehicle-injected *Cib2*^*KO*^ controls (**Fig. 6b**).

**Figure 6:**
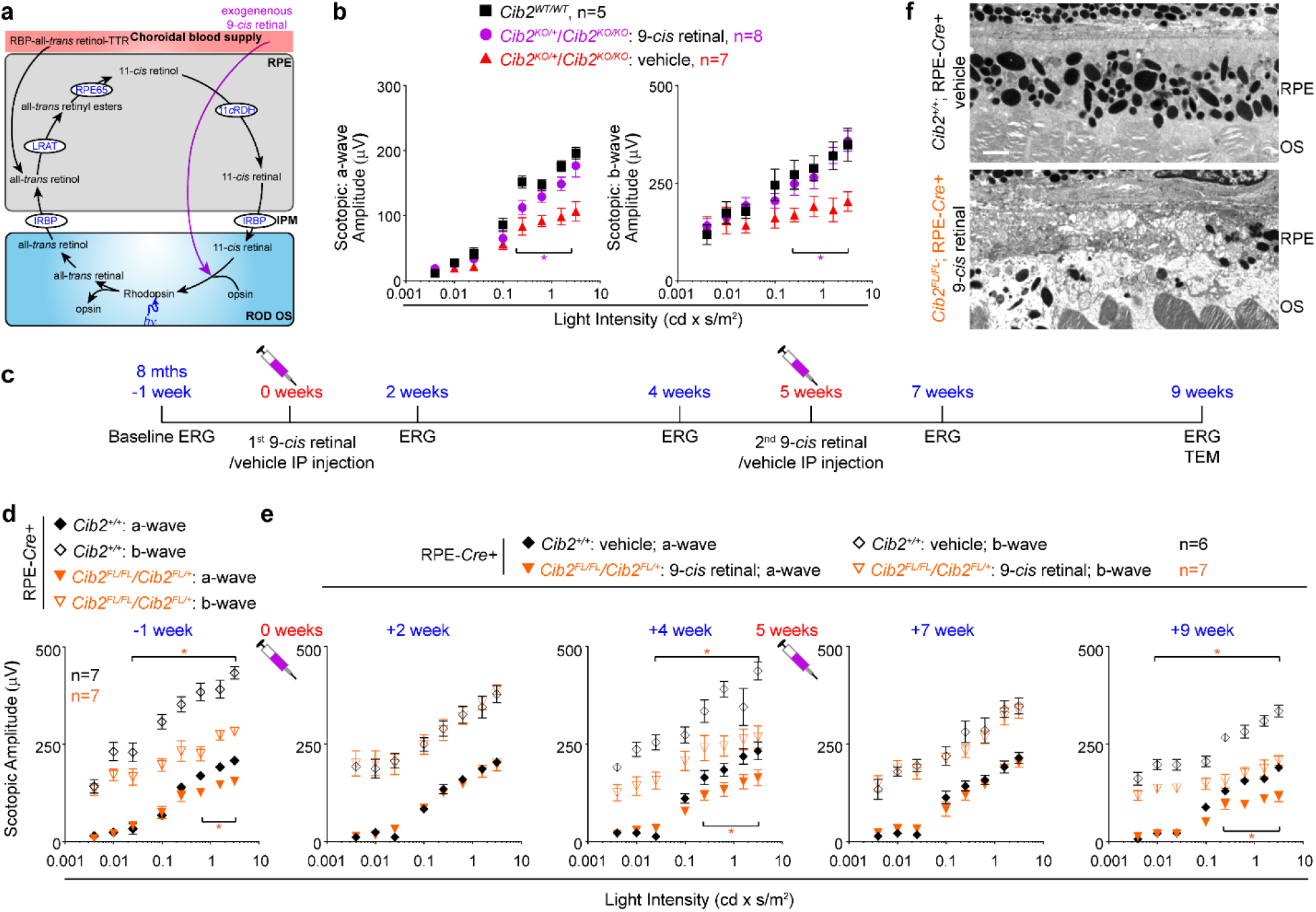
PR function but not RPE pathology can be rescued by exogenous retinoid. **a** Schematic of the visual cycle and exogenous 9-*cis* retinal treatment. **b** Eight-to-nine month old *Cib2* mutant mice (*Cib2*^*KO/+*^ and *Cib2*^*KO/KO*^) were either intraperitoneally injected with vehicle (*n*=7) or 0.25 mg 9-*cis* retinal (*n*=8), dark adapted overnight and ERG performed next day. Both a-wave (*left* panel) and b-wave (*right* panel) amplitudes were higher in mutant mice injected with 9-*cis* retinal than vehicle-injected mice. For comparison untreated WT control mice (black squares, *n*=5) are replotted from **Fig. 1c**, *bottom* panels. **c** Schematic of treatment paradigm in RPE-specific *Cib2*^*KO*^ mice injected with 9-*cis* retinal and control mice injected with vehicle. **d** Baseline ERG for indicated genotype are reproduced from **Fig 2a**, *bottom* panels. **e** ERG amplitudes for indicated genotype after specific treatment as outlined in **c** (vehicle injected; *n*=6, 9-*cis* retinal injected *n*=7). **f** TEM micrographs for indicated genotype at the end of the treatment paradigm as outlined in **c** (*n*=3/genotype). *Scale bar*, 2 µm. Unpaired two-tailed *t* test, *p* value of at least <0.05 (*).

Next, to further assess the potential clinical translatability of exogenous retinoids, we employed an extended treatment paradigm following injection of 9-*cis* retinal in RPE-specific *Cib2*^*KO*^ mice (**Fig. 6c**). We performed baseline ERGs on eight-month-old uninjected mice and found significant differences in ERG amplitudes in mutant mice compared to RPE-*Cre+* control mice (**Fig. 6d**). Two weeks after the first injection, we found significantly improved ERG amplitudes for RPE-specific mutant mice injected with 9-*cis* retinal. The ERG amplitudes of 9-*cis* retinal injected mice were essentially indistinguishable from those of vehicle-injected control mice. However, the rescue effect of exogenous retinoid was short-lived. By four weeks post-injection, the ERG amplitudes of 9-*cis* retinal injected RPE-specific mutant mice were substantially lower than control mice. To assess the medium-term efficacy of exogenous retinoid therapy, we extended the paradigm further by one month. Again, we saw an improvement in ERG amplitudes for mutant mice injected with 9-*cis* retinal two weeks after the second injection, but by four weeks, ERG amplitudes fell back to baseline levels (**Fig. 6e**). This suggested that short-term but reproducible improvement in ERG amplitudes are possible with this treatment. At the conclusion of treatment, TEM analysis of the RPE revealed vacuoles and/or sub-RPE deposits present in all three 9-*cis* retinal-treated RPE-specific *Cib2*^*KO*^ mice, but not in vehicle-treated RPE-*Cre+* control mice (1 of 3) (**Fig. 6f**). The results strongly suggest that repeated exogenous retinoid treatment can rescue photoreceptor function temporarily yet reproducibly in *Cib2* deficient mice by bypassing the RPE. Although, (as expected) this does not improve RPE pathophysiology.

## DISCUSSION

This study highlights the crucial role of CIB2 in the regulation of phagolysosomal digestion of OS and development of age-related phenotype in mice. CIB2 belongs to a family of proteins that includes four members (CIB1-CIB4). CIB proteins contain Ca^2+^/Mg^2+^ binding EF-hand domains, and binding of these ions leads to conformational change and downstream effects, similar to the well-studied protein, calmodulin ^34,35^. In recent years, the various roles of CIB2 have been elucidated in multiple organs development or function, however, the precise tissue-specific mechanisms remain unclear. For instance, *CIB2* is down-regulated in ovarian cancer and associated with poor prognosis in ovarian cancer patients ^36^. Previously, we and others reported on the essential function of CIB2 in the inner ear development and mechanotransduction of sound signal ^16,37,38^. Furthermore, *Cib2* deficient mice showed small, but not statistically significant difference in ERG amplitude between heterozygous and homozygous mice, leading the authors to conclude that CIB2-deficient mice exhibit no retinal phenotype ^37^. However, the main caveats of this study were use of only one intensity of light for retinal functional evaluation and lack of instrumental WT controls ^37^.

Further, we used three different mutant mouse strains and found that both a complete loss (homozygous) and haploinsufficiency (heterozygous) of CIB2, specifically in RPE, leads to attenuated ERG amplitudes and development of age-related phenotype encapsulating several dry AMD features. The mutant mice from our study had fewer lysosomes, lysosomal, and autophagy proteins and defects in phagolysosomal clearance of OS. As OS phagosomes contain ∼50% lipid, chronically delayed OS clearance causes accumulation of lipids. Confirming this trend, we found double the amount of neutral lipid droplets in the RPE in aged *Cib2*^*KO*^ mice with gradual accumulation of un-cleared lipids and proteins. Since RPE cells are post-mitotic these insults accumulate over time, resulting in an age dependent RPE pathology and PR dysfunction. Earlier, genetic deletion of genes involved in clearance of ingested OS, specifically in the RPE, was found to lead to secondary photoreceptor functional defects ^2,9,18^. Further, the RPE presents the largest source of electrical resistance for *in vivo* ERGs ^39^, and hence the ERG changes we observed were likely caused by RPE dysfunction and subsequent pathology. Our PR- and RPE-specific *Cib2* knockout data further support this conclusion.

The RPE-mediated OS non-inflammatory clearance process requires LC3-associated phagocytosis. Deficiency in LAP has been associated with various disorders such as SLE, multiple sclerosis, atherosclerosis, and AMD, amongst others ^40^. It has proven quite challenging to study AMD, since mouse models of genetically validated targets do not recapitulate the human phenotype. Here, we show that global or RPE-specific lack of CIB2 recapitulates many aspects of the human AMD pathophysiology including sub-RPE deposits, and accumulation of APOE, C3, and aβ. Further, we found increased lipid droplets and esterified cholesterol, which is associated with drusen in humans and sub-RPE deposits in mouse models. A recent study using RPE-specific LAMP2 knockout mouse showed faulty processing of ingested OS, lipid accumulation, and similar RPE deposit protein composition ^28^ as compared to this current study. A growing amount of scientific literature, in which phagolysosomal digestion is impaired due to genetic loss of specific proteins in the RPE, also shows similar phenotype as our model ^2,9,18,28^, further supporting the speculation that dysregulation of OS processing may be a causal factor in AMD pathophysiology. Additionally, the mutant *Cib2* mouse may serve as model to study AMD and for translational efforts to halt AMD. Furthermore, there are many mechanistic and molecular similarities between LAP and canonical autophagy. This would be an interesting area of future investigations to assess if CIB2 also participates in the regulation of conventional autophagy.

Developing translational models to study AMD progression is imperative to enable evaluation of rational treatments for AMD. Patients with dry AMD in particular have little to no treatment options, although the AREDS (Age-Related Eye Disease Study) nutritional supplementation formula, with carotenoids, vitamins, zinc, and copper, have been shown to have a modest effect on progression to neovascular AMD. Bypassing the RPE visual cycle by supplying exogenous retinoids has been found to effectively serve as a useful therapeutic approach for certain retinal disorders ^41,42^. The most common methodology is using a 9-*cis* retinoid analog that is more stable than the physiological 11-*cis* retinal chromophore. Upon entering the PR, 9-*cis* retinal can bind with opsin to create functional visual pigment. However, unbound retinals are quite cytotoxic and quickly detoxified by the liver – an issue that has been circumvented by using the less toxic oral 9-*cis* retinyl acetate to treat inherited blindness of childhood caused by RPE65 and LRAT mutations ^43^. We have found that *Cib2*^*KO*^ RPE contain abundant large lipid-filled vacuoles and sub-retinal deposits, which may impede RPE transport mechanisms including those that deliver retinoids back to the PR, leading to reduced ERG amplitudes, even without gross PR damage. Supporting this idea, we found that providing exogenous retinoids improved ERGs in two different treatment paradigms in two different mouse models, consistent with a small study in humans ^44^. In summary, our findings reveal a new mechanistic model of AMD elicited by a CIB2 deficiency, specifically in the RPE, resulting in pathogenic impairment of LAP and secondarily visual function. Further, we have also provided pre-clinical proof-of-concept for use of retinoids to treat AMD.

## METHODS

### Animals, tissue harvest, and processing

The ARRIVE guidelines for reporting animal research were used for procedures involving animals. All animal studies were conducted according to the ARVO *Statement for the Use of Animals in Ophthalmic and Vision Research* and the National Institutes of Health *Guide for the Care and Use of Laboratory Animals*, and after approval from the IACUC (Institutional Animal Care and Use Committees) of the University of Maryland School of Medicine. *Cib2*^*tm1a*^ mice (designated *CIB2*^*KO*^) were purchased from EUCOMM and have been validated previously ^16,37^. To generate floxed allele (*Cib2*^*FL*^), *Cib2*^*tm1a*^ were crossed ROSA26::FlPe knock in mice (Jax lab, stock#003946), which resulted in the removal of LacZ-neomycin selection cassette by Flp-FRT recombination. RPE-specific *Cre* expressing mice were described earlier ^20^, and were generously shared with us by Drs. Sheldon Miller, National Eye Institute and Joshua Duniaef from University of Pennsylvania. Previous research pointed out that these mice may develop *Cre*-mediated age-dependent RPE pathology ^45^, hence for all experiments we used *Cib2*^*+/+*^; RPE-*Cre*+ mice as controls. Rod photoreceptor-specific *Cre*-expressing mice ^19^ were purchased from Jackson Labs. Previous studies and our own studies have not shown any Cre-mediated toxicity in this line, hence for controls we used a mix of *Cib2*^*FL/FL*^ or *Cib2*^*FL/+*^; *Cre-*, and *Cib2*^*+/+*^; *Cre+* mice. Mice were housed under a strict 12:12 h light on/off cycle and fed standard mouse diet (after weaning) and water *ad libitum*. Mice were euthanized by CO_2_ asphyxiation followed by cervical dislocation before immediate eye enucleation. Eyes were immersion-fixed in 2% PFA + 2% glutaraldehyde for morphometry and TEM.

RPE flat mounts were generated by first removing the anterior segment and lens from an eye, and then removing the retina carefully in Hank’s balanced salt solution (HBSS). After fixing the eyecup in 4% PFA in PBS for 30 mins to 1 hr, radial cuts were made to flatten the eyecup, and antibody/dye labeling was performed for fluorescence microscopy. For immunoblots, RPE/choroid tissues were snap-frozen on dry ice before storing at −80°C before lysis. Uncropped blots are presented in **supplementary Fig. 7**. For retinoid analysis, mice were dark-adapted overnight, and dissections were performed under dim red light.

### Morphometric analysis of the outer retina

Plastic sections were prepared as previously described ^46^. Eyes were hemisected from the pupil through the optic nerve and the eyecups harvested, dehydrated in ascending ethanol concentrations, infiltrated, and embedded in JB-4 Plus resin (Ted Pella, Redding, CA). Sections 4 µm thick were cut on a Leica EMUC6 Ultramicrotome (Leica Microsystems, Buffalo Grove, IL), mounted on slides, dried, and stained with 1% cresyl violet (Sigma, St. Louis, MO). Sections were examined at 40X or 100X using a NIKON EFD-3 Episcopic-Fluorescence microscope (Nikon Inc., Melville, NY). Photomicrographs were taken using a Moticam 2500 high-resolution camera (Motic, British Columbia, Canada) and brightness and contrast adjusted as needed using Adobe Photoshop (Adobe Systems, San Jose, CA).

To measure the thicknesses of outer retina layers, a vertical line was drawn from the outer plexiform layer (OPL) to the RPE, and the OPL, outer nuclear layer (ONL), PR OS, and distances of inner segment (IS) edges measured individually from this line using Moticam Image Plus 2.0 Software (Motic China Group Co., Ltd., Xiamen, China), in 10 sections per eye, and overall means calculated. Retinal thickness was measured without knowledge of the genotype by a masked observer (PAS).

### Transmission electron microscopy

Sections for EM were prepared as previously described ^46^. Briefly, hemisected eyecups were rinsed in buffer and postfixed in 2% osmium tetroxide and 1.5% potassium ferrocyanide in dH_2_O for 2 hr, dehydrated in a graded ethanol series, and embedded in Epon-Araldite (Electron Microscopy Sciences, Hatfield, PA). Semi-thin sections (4 μm) were cut and stained with 1% cresyl violet. Ultra-thin sections (90 nm) were cut on an ultramicrotome (Ultracut E 701704, Reichert-Jung, Buffalo, NY) using a diamond knife (Micro Star Technologies, Inc., Huntsville, TX), collected on copper grids, counterstained with 4% methanolic uranyl acetate (Electron Microscopy Sciences, Hatfield, PA), and outer retinal morphology examined using a transmission electron microscope (TEM; Model 300: Phillips, Eindhoven, The Netherlands). Photomicrographs were captured with a digital camera (15-megapixel digital camera, Scientific Instruments and Applications, Duluth, GA) and Maxim DL Version 5 software (Diffraction Limited, Ottawa, Canada).

### Electroretinography

Electroretinograms (ERG) were recorded as earlier described ^18^ with modifications. Mice, overnight dark-adapted, were anesthetized with ketamine-xylazine (100 mg/kg and 10 mg/kg, respectively) and pupils were dilated with 1% Tropicamide. A gold wire lens electrode was placed on the cornea, a platinum reference electrode in the mouth, and a ground electrode on the tail. To assess rod-driven responses, increasing scotopic stimuli were presented sequentially (0.003962233 to 3.147314 cd x s/m2) at 5-60 sec intervals using UTAS BigShot (LKC Technologies, Gaithersburg, MD). At least 3 waveforms per intensity were averaged. For cone function evaluation, photopic responses to a single bright flash (3.15 cd x s/m2) under a steady rod-suppressing field of cd x s/m2. Waves were analyzed using EM for Windows software (LKC Technologies).

### Cell culture, drug treatment, synchronized cell culture phagocytosis assay

Immortalized RPE-J cells derived from rat (ATCC, Manassas, VA) were maintained at 32°C and 8% CO_2_ in 4% FBS/DMEM supplemented with 1x penicillin/streptomycin. HEK-293T, COS-7, LAM-621, and *TSC2*^*KO/KO*^ *p53*^*KO/KO*^ MEF cells were maintained at 37°C/ 5% CO2 in 10% FBS-DMEM supplemented with 1x penicillin/streptomycin. OS were purified from fresh porcine eyes harvested within 24 hrs from Sierra For Medical Science (Whittier, CA) using an established protocol ^47^, modifying only the preparation of sucrose density gradient tubes. 24 hrs before OS purification, sucrose step gradients (20%-60% in 10% increments) were prepared in ultracentrifugation tubes and frozen at −20°C at an approximately 45° angle. On the day of purification, a continuous gradient was formed by thawing at room temperature at this angle. Pulse-chase experiments were performed as we described ^18^. RPE-J cells transiently overexpressing AcGFP or CIB2-IRES-AcGFP were challenged with ∼10 OS/cell in serum-free DMEM with 1 μM MFG-E8 for 1 hr at 20°C (pulse, POS binding only). After washing unbound POS with PBS, cells were chased for indicated times at 32°C with 5% FBS-supplemented DMEM. At the end of the pulse or chase period, cells were rinsed 3x with PBS before lysis in RIPA buffer with a solution containing 1x protease inhibitors, 1 mM Na orthovanadate, 10 mM Na glycerophosphate, and 10 mM NaF, and stored at −80°C until immunoblotting.

### X-gal staining, lipid droplet, and indirect lysosome labeling

For X-gal staining, *Cib2*^*WT/WT*^ (control) and *Cib2*^*KO/+*^ enucleated eyes were fixed for 1 hr in 0.5% glutaraldehyde (0.5%)+NP-40 (0.02%) diluted in PBS, followed by washing with PBS 2x for 5 min. Eyes were stained overnight at 37°C in tubes covered with aluminum foil with PBS, in a solution of 1 mg/ml X-Gal, 5 mM K_3_Fe(CN)_6_, 5 mM K_4_Fe(CN)_6_, 2 mM MgCl_2_, 0.02% NP-40, and 0.01% sodium deoxycholate. The next day, the eyes were washed once for 5 min in PBS followed by post-fixation with 2% PFA – 2% glutaraldehyde in PBS for at least 1 hr at RT. An opening was made in the lens region to allow OCT penetration and embedding. 20-30 µm thick sections were cut on a Cryotome (Leica, Germany).

Phagosome counts in whole mounts of RPE were performed using Ret-P1 antibody ^18^. Whole-mount RPE/choroid preparations were live-stained with 1 µM BODIPY-Pepstatin A or 0.1 mg/ml BODIPY-493/503 in DMEM at 37°C for 30 min followed by fixation with 4% PFA.

60-70% confluent COS-7 cells or RPE-J cells were fixed with 4% PFA for 15 min at RT and permeabilized with 0.2 % Triton X-100 in PBS (15 min), followed by blocking with 10% normal goat serum (NGS) in PBS for at least 30 min at RT. Primary antibodies were diluted in 3% NGS-PBS and incubated overnight at 4°C, followed by the incubations with the indicated goat secondary antibodies. Phalloidin-647 or rhodamine-phalloidin was added at dilutions of 1:1000 (COS-7) or 1:200 (RPE-J) during secondary antibody incubation to stain F-actin. A Zeiss 710 laser scanning confocal microscopy system was used for image acquisition, with step size of 0.5 µm. Fiji (ImageJ) ^48^ was used to process images, and average numbers of phagosomes per 100 µm^2^ of retina were calculated [detailed in ref ^8^]. For counting area of undigested material or mitochondria as control (**Fig. 2c, d**), outlines were drawn in Fiji on TEM images from indicated genotypes of mice sacrificed 8 hr after light onset, and areas calculated using Fiji functions.

### Filipin and Oil red O staining

Ten micron cryosections were prepared using standard procedures and filipin staining for esterified cholesterol (EC) and unesterified cholesterol (UC) was performed ^28^. For oil Red O staining, cryosections were dipped in 60% isopropanol for 5 seconds, followed by incubation for 15 mins at RT in working solution of Oil Red O (3 parts:2 parts::0.5% Oil Red O solution in isopropanol:water, the working solution was mixed and allowed to stand for 10 mins before filtration with vacuum filtration system containing 0.22 µm filter). The slides were then dipped in 60% isopropanol 5 times to remove excess stain and dipped in distilled water 10 times. The imaging was performed with a 100x objective.

### Transfection and immunoblotting

We optimized the following protocol for 24 wells RPE-J cells transfection using Lipofectamine-2000: Lipofectamine 2000:DNA (2-4 µg plasmid/well)::3:1 was prepared in 100 µl Optimem. Although, the very high plasmid concentration leads to toxicity for most other cell types but works well for RPE-J cells. Cells were used for transfection at ∼70% confluence. The medium was replaced with 400 µl Optimem/well (adding Optimem instead of 2% FBS-DMEM is important for transfecting RPE-J cells). The Lipofectamine/plasmid mix was added dropwise. 24 hrs later the medium was changed to complete medium (4% FBS-DMEM), and cells were allowed to grow/express plasmid for further 24 hrs before fixing or used for pulse-chase experiments, followed by immunoblotting. This method reliably and reproducibly gave us >60% transfection.

### Retinoid extraction and analysis

All procedures were performed under red safelights. For analysis of mouse ocular retinoids, mouse eye tissues (retinae or eyecups) were homogenized in 1 ml of freshly made hydroxylamine buffer (50 mM MOPS, 10 mM NH2OH, pH 6.5). After addition of 1 ml ethanol, samples were mixed, then incubated for 30 min in the dark at RT. Retinoids were extracted into hexane (2 × 4 ml) then solvent was evaporated under a gentle stream of argon at 37°C. The dried samples were re-dissolved in 50 μl mobile phase for analysis. Retinaloxime ^49^ and retinyl palmitate (Sigma Aldrich, Saint Louis, MO) standards, and samples were separated on a 5 μm LiChrospher Si-60 (ES Industries, West Berlin, NJ) normal-phase column on a Waters H-Class Acquity UPLC (Waters Corp., Milford, MA, USA) in hexane mobile phase containing ethyl acetate (11.2%):dioxane (2.0%):octanol (1.4%; (Landers & Olson, 1988)) at a flow rate of 1.5 ml/min. Absorbance was monitored at 325 nm for retinyl esters and 350 nm for retinaloximes. Peak areas were integrated and quantified using calibration curves based on external standards. Data were analyzed using Empower 3 software (Waters Corp.).

### A2E extraction and analysis

Pairs of mouse eyes from individual mice were homogenized in 975 µl H_2_O and 2.5 ml methanol-glacial acetic acid (98:2), followed by addition of 1.125 ml chloroform to get a single-phase solution. The solution was thoroughly mixed and centrifuged for 5 min at 13,000 rpm to pellet proteins. Then 1.225 mlL of chloroform and 1.225 ml of H_2_O were added, the tube was mixed for 1 min to separate into two phases and the lower organic phase containing A2E was collected. The solvent was evaporated, and the residue dissolved in acetonitrile (ACN) for analysis. A2E was analyzed by reversed phase HPLC on a Waters Acquity H-Class UPLC system equipped with an Acquity UPLC BEH C18 column (1.7 μm particles; 2.1 x 50 mm) using isocratic elution with 0.1% trifluoroacetic acid in 20% H_2_O/80% ACN at 1.2 ml/min. Specific wavelength detection was used for monitoring A2E (430 nm), and the quantities of A2E were determined from a calibration curve developed using a synthetic A2E standard. Data were analyzed using Empower 3 software (Waters Corp.).

### Data analysis

For ERG analysis, four to eight animals were used per time point/genotype/treatment. For *in vivo* experiments, at least three mouse eyes (from different animals) were used for immunoblots, at least three images were averaged per eye from the three eyes/indicated genotype/time point for confocal microscopy. For cell culture (*in vitro*) experiments, at least three independent experiments or two independent experiments for co-IP studies were performed. One-way ANOVA with Tukey’s post-hoc test or a student’s *t*-test was used to compare control sample to test samples, with data presented as mean ± SEM. Differences with p < 0.05 were considered significant. Data were analyzed using GraphPad Prism (GraphPad Software, Inc., La Jolla, CA).

| REAGENT or RESOURCE | SOURCE | IDENTIFIER |
| --- | --- | --- |
| **ANTIBODIES (Anti)** |  |  |
| CIB2 (clone 59, homemade) | This paper |  |
| Actin (clone 13E5) | Cell Signaling Technology | # 4970S |
| β-Tubulin (clone 9F3) | Cell Signaling Technology | # 2128S |
| HA.11 Epitope Tag | Biolegend | # 901501 |
| Cathepsin D | R & D systems | AF1029 |
| ATG5; N-terminal | Sigma | A0856 |
| LAMP1 | Sigma | L1418 |
| Rhodopsin Antibody, clone RET-P1 | EMD Millipore | MAB5316 |
| Opsin (clone B6-30); do not boil sample for immunoblot | Novus Biologicals | NBP2-25160 |
| Apolipoprotein E | Abcam | ab183596 |
| C3 | Abcam | ab97462 |
| Beta-amyloid (d3d2n) | Cell Signaling Technology | CST-15126T |
| LAMP2/ Cd107b | Bio-Rad | RAD-AHP1634 |
| **REAGENTS** |  |  |
| ECL™ Prime Western Blotting System | Thermo Fisher | 32106 |
| Lipofectamine™ 2000 Transfection Reagent | Thermo Fisher | 11668019 |
| In-Fusion® HD Cloning Plus | Clontech | 638911 |
| BODIPY 493/503 | Thermo Fisher | D3922 |
| Pepstatin A, BODIPY™ FL Conjugate | Thermo Fisher | P12271 |
| X-Gal | Sigma-Aldrich | 3117073001 |
| 9-*cis* retinal | Sigma-Aldrich | R5754-100MG |
| Filipin Iii | Sigma-Aldrich | SIG-F4767- |
| Oil red O | Sigma-Aldrich | O0625 |
| **Recombinant Proteins** |  |  |
| Recombinant Human MFG-E8 Protein | R & D systems | 2767-MF-050 |
| **Experimental Models: Cell Lines** |  |  |
| RPE-J | ATCC | # CRL-2240 |
| **Recombinant DNA** |  |  |
| pAAV-CIB2-IRES-AcGFP | This paper |  |
| pAAV-AcGFP | Grousbeck Gene Therapy Center |  |
| **Software and Algorithms** |  |  |
| EM for Windows | LKC |  |
| Prism 6.0 | GraphPad, La Jolla, CA |  |
| Adobe Suite | Adobe, CA |  |
| Fiji | https://fiji.sc/ |  |
| **Mouse strains** |  |  |
| *Cib2<sup>tm1a(EUCOMM)Wtsi</sup>* mice | EUCOMM | EM:05417 |
| *VMD2*-Cre mice | Gift from Drs. Sheldon Miller and Joshua Dunaief | Iacovelli et al., 2011 |
| *Opsin-iCre75* | Jax | Stock No: 015850 |
| ROSA26::FIPe knock in mice | Jax | Stock No: 003946 |

### Contact for reagent and resource sharing

Further information and requests for resources and reagents should be directed to and will be fulfilled by the Lead Contact, Zubair Ahmed, PhD.

## Contributions

S.S. and Z.M.A. designed and conceived the project. S.S., P.A.S, A.P.J.G., T.D. carried out experimental work and analyzed data. S.R., T.M.R., and Z.M.A. supervised the experiments. S.S. and ZMA wrote the manuscript. All authors read, edited, and approved the manuscript.

## Acknowledgments

We thank Drs. M. Johnson, M. Matthews, and Ms. D. Gomes for technical assistance; Drs. S. Miller, and J. Duniaef for mouse strains; and the UMSOM core facility for access to the Zeiss-710 confocal microscope. We also thank Drs. T. Friedman, R. Hertzano, C. Mueller, G. Frolenkov and Ms. M. Ahmed for critically reading the manuscript. This work was supported by NIDCD/NIH grants R01DC012564, R01DC016295 (Z.M.A.), R56DC011803 (S.R.), and partly by the Loris Rich Postdoctoral fellowship, International Retinal Research Foundation, AL, USA (S.S.), and Research to Prevent Blindness Award (P.A.S., Z.M.A.).

